# O-GlcNAc Homeostasis Controls Cell Fate Decisions During Hematopoiesis

**DOI:** 10.1101/337881

**Authors:** Zhen Zhang, Matt P. Parker, Stefan Graw, Lesya V. Novikova, Halyna Fedosyuk, Joseph D. Fontes, Devin C. Koestler, Kenneth R. Peterson, Chad Slawson

## Abstract

The addition of O-GlcNAc (a single β-D-N-acetylglucosamine sugar at serine and threonine residues) by O-GlcNAc transferase (OGT) and removal by O-GlcNAcase (OGA) maintains homeostatic levels of O-GlcNAc. We investigated the role of O-GlcNAc homeostasis in hematopoiesis utilizing G1E-ER4 cells carrying a GATA-1 transcription factor fused to the estrogen receptor (GATA-1ER) that undergo erythropoiesis following the addition of β-estradiol (E2) and myeloid leukemia cells that differentiate into neutrophils in the presence of all-trans retinoic acid. During G1E-ER4 differentiation, a decrease in overall O-GlcNAc levels and an increase in GATA-1 interactions with OGT and OGA were observed. Transcriptome analysis on G1E-ER4 cells differentiated in the presence of Thiamet-G (TMG), an OGA inhibitor, identified expression changes in 433 GATA-1 target genes. Chromatin immunoprecipitation demonstrated that the occupancy of GATA-1, OGT, and OGA at *Laptm5* gene GATA site was decreased with TMG. Myeloid leukemia cells showed a decline in O-GlcNAc levels during differentiation and TMG reduced the expression of genes involved in differentiation. Sustained treatment with TMG in G1E-ER4 cells prior to differentiation caused a reduction of hemoglobin positive cells during differentiation. Our results show that alterations in O-GlcNAc homeostasis disrupt transcriptional programs causing differentiation errors suggesting a vital role of O-GlcNAcylation in control of cell fate.

## Introduction

Hematopoiesis starts during embryonic development and occurs throughout adulthood to produce and replenish the blood system. Hematopoiesis leads to distinct cellular linages by the coordination of defined gene expression programs. The spatiotemporal coordination of gene expression is encoded in the genome and read by lineage-specific transcription factors to active or inactive genes (Goode et al. 2016). The molecular mechanisms by which transcriptional programs are established and maintained in hematopoietic cells is broadly understood but how post-translational modifications (PTMs) refine these transcriptional programs in response to changes in the environment is uncertain. Understanding how the cellular environment controls hematopoiesis by PTMs has important implications in our understanding of the modulation of linage specific gene pathways and the potential development of novel therapies to treat blood-related disorders.

Erythropoiesis, for example, is the process by which hematopoietic progenitors proliferate and differentiate into red blood cells. Erythropoiesis is regulated by numerous transcription factors that orchestrate cellular differentiation. GATA-1, a transcription factor bearing a dual-zinc finger that recognizes a WGATAR DNA motif, is a key transcription factor that coordinates erythroid differentiation gene programs (Yamamoto et al. 1990; Pevny et al. 1991; Orkin 1992; Fujiwara et al. 1996). During erythroid differentiation, many GATA-1 target genes are activated or repressed (Welch et al. 2004; Cheng et al. 2009; Wu et al. 2011; Kingsley et al. 2013). The co-occupancy of GATA-1 with erythroid transcription factors, activators/repressors, and co-activators/corepressors can determine the GATA-1 target gene transcription status to some extent (Cheng et al. 2009; Tripic et al. 2009; Yu et al. 2009; Soler et al. 2010). Epigenetic modifications of histones and variation of WGATAR adjacent motifs also play a role (Cheng et al. 2009; Fujiwara et al. 2009; Zhang et al. 2009; Bresnick et al. 2010; Wu et al. 2011). However, these mechanisms do not explain the full breadth of GATA-1 gene regulation.

Potentially, PTMs of transcription complexes during hematopoiesis could regulate linage specific transcription factor activity by altering protein-protein interaction with transcriptional co-activators/co-repressors, or by modifying chromatin structure (Boyes et al. 1998; Lamonica et al. 2006; Lee et al. 2009; Kadri et al. 2015). Recently, we demonstrated that O-GlcNAcylation modulates GATA-1-mediated repression of the ^A^γ-globin promoter (Zhang et al. 2016). O-linked β-N-acetylglucosamine (O-GlcNAc) is the modification of serine and threonine residues by β-D-N-acetylglucosamine. This sugar moiety is added and removed from serine and threonine residues by the O-GlcNAc processing enzymes, O-GlcNAc transferase (OGT) and O-GlcNAcase (OGA), respectively (Hart 2014). Dynamic O-GlcNAc cycling is critical for the proper regulation of gene transcription. O-GlcNAc is one of the histone code marks (Sakabe et al. 2010; Gambetta and Muller 2015) and modulates other epigenetic changes on chromatin such as DNA methylation (Chen et al. 2013; Deplus et al. 2013), histone acetylation (Yang et al. 2002), and histone methylation (Deplus et al. 2013; Chu et al. 2014). For example, TET (ten-eleven translocation) proteins and HCF-1 (host cell factor 1) co-localize with OGT at gene promoters associated with activating histone marks to promote target gene transcription (Dey et al. 2012; Chen et al. 2013; Deplus et al. 2013; Gambetta and Muller 2015). OGT also interacts with mSin3A or components of the polycomb repressive complex (PRC) 1 and 2 to mediate transcription repression by forming stable repressor complexes (Yang et al. 2002; Chu et al. 2014; Maury et al. 2015). OGT and OGA regulate preinitiation complex (PIC) formation and RNA Polymerase II (Pol II) activity (Ranuncolo et al. 2012; Lewis et al. 2016; Resto et al. 2016).

OGT and OGA interact with the GATA-1-FOG-1-Mi2β repressor complex at the ^A^γ-globin promoter, and O-GlcNAcylation of chromatin remodeler Mi2β modulates GATA-1-FOG-1-Mi2β repressor complex formation to negatively regulate ^A^γ-globin transcription (Zhang et al. 2016). Since O-GlcNAc cycling on chromatin modifiers is critical to modulate gene transcription and normal development (Howerton et al. 2013; Olivier-Van Stichelen and Hanover 2015), potentially, O-GlcNAc could regulate hematopoietic cell differentiation and development. Overall O-GlcNAc levels decrease during embryonic stem cell (ESC) differentiation into neurons (Speakman et al. 2014), epidermal keratinocytes (Sohn et al. 2014), cardiomyocytes (Kim et al. 2009), and myoblasts (Ogawa et al. 2012). In addition, O-GlcNAc levels decrease in K562 erythroleukemia cells following treatments that induce terminal differentiation (Zhang et al. 2016); however, the overall O-GlcNAc levels increase during differentiation of mouse chondrocyte (Andres-Bergos et al. 2012) and osteoblast cells (Koyama and Kamemura 2015). GATA-1 is the master regulatory transcription factor for erythropoiesis and our previous work established a role for O-GlcNAc cycling in the modulation of GATA-1-FOG-1-Mi2β repressor complex. We hypothesize that O-GlcNAc homeostasis contributes regulates gene expression programs during hematopoietic cell differentiation.

To test this hypothesis, we used well-established erythropoiesis and myeloid cell systems. The G1E-ER4 cell line that stably expresses GATA-1-ER, a fusion product of GATA-1 and human estrogen receptor ligand binding domain (Welch et al. 2004), is a hematopoietic cell line derived from murine G1E (GATA-1^−^ erythroid) cells. When GATA-1 activity is restored by β-estradiol (E2), G1E-ER4 cells undergo red blood cell differentiation and recapitulate the developmental stages from late BFU-E (burst forming unit-erythroid) to basophilic erythroblasts (Welch et al. 2004; Grass et al. 2006). The NB4 and HL60 myeloid leukemia cell lines differentiate into neutrophil-like cells with the addition of all-trans retinoic acid (Drach et al. 1993). In this work, we demonstrate that: 1) hematopoietic cell differentiation caused a marked reduction in total O-GlcNAc levels; 2) interactions between GATA-1-ER, OGT, and OGA were enhanced after β-estradiol induced differentiation; 3) perturbation of O-GlcNAc cycling led to 1,173 differentially expressed genes, including 47 erythroid-specific GATA-1 target genes; 4) OGA inhibition leads to an amplification of GATA-1 activation or repression; 5) TMG treatment impaired neutrophil-like cells differentiation and 6) prolonged TMG treatment severely disrupted erythropoiesis. As a result of these findings, we conclude that O-GlcNAcylation is a mechanism to modulate linage-specific transcriptional programs controlling hematopoietic differentiation.

## Results

### O-GlcNAc levels decrease during erythropoiesis

To determine if O-GlcNAcylation is involved in erythroid differentiation, we treated GATA-1-ER cells with E2 to trigger erythroid differentiation. Expression of GATA-1-ER increased as previously reported (Lamonica et al. 2011; Manavathi et al. 2012); *Gata2* transcription was repressed by GATA-1, resulting in the decrease of GAtA-2 protein level (Bresnick et al. 2010). Expression of Friend of GATA-1 (FOG-1) also increased as previously reported (Welch et al. 2004) (**Figure 1a**). Interestingly, the total cellular O-GlcNAc levels were dramatically decreased after E2 treatment in G1E-ER4 cells. We found that the O-GlcNAc levels continued to decrease as differentiation proceeded, with 30 hrs being the lowest compared to other time points. Not surprisingly, the decreased O-GlcNAc levels were associated with a reduction in OGT protein levels and an increase in OGA protein levels, as these enzymes are reciprocally regulated to maintain appropriate levels of O-GlcNAcylation in cells (**Figure 1a**). However, the large decrease in O-GlcNAc levels after E2 treatment suggests that the targeting of OGT and OGA to specific substrates might also be altered during differentiation. Importantly, O-GlcNAc levels, OGT, and OGA expression in G1E negative control cells did not change over time (**Figure 1a**). Together, these data indicate that the O-GlcNAc levels decrease during erythropoiesis, and the decline was a direct result of restoration of GATA-1 activity.

**Figure 1.**
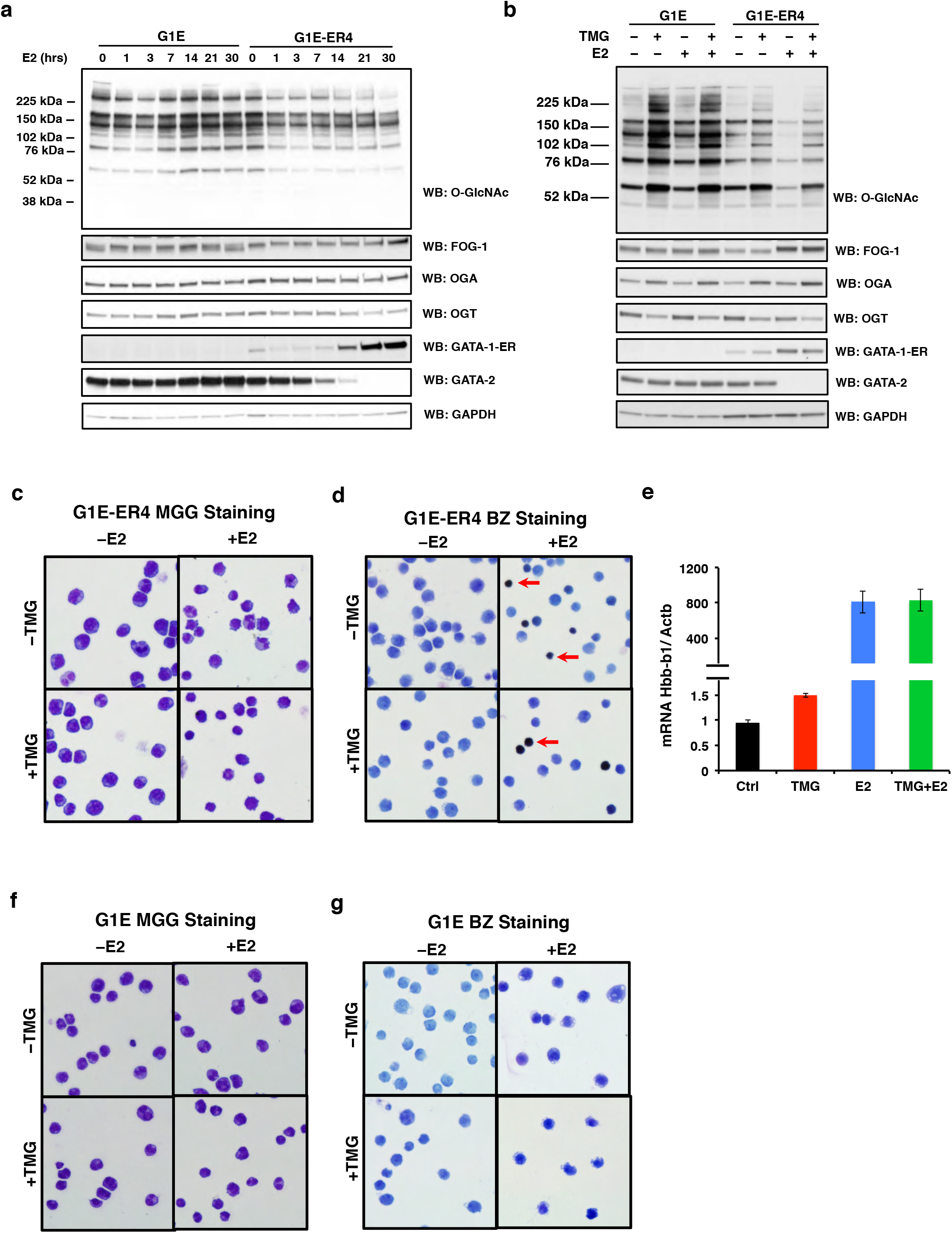
O-GlcNAc levels decrease following restoration of GATA-1 activity. G1E and G1E-ER4 cells were treated with TMG (OGA inhibitor) and/or E2 for 30 hrs. Cells were harvested (**a**) following a time course or (**b**) at 30 hrs and total cell lysates were subjected to immunoblotting. G1E-ER4 (**c** and **d**) and G1E (**f** and **g**) cells were subjected to May-Grunwald Giemsa (MGG) staining (c and f) for cell morphology and benzidine staining (BZ) (**d** and **g**) for hemoglobin content. Red arrow shows hemoglobin positive cells. **e**. Transcription level of β^major^-globin gene (*Hbb-b1*) was measured by qPCR and β-actin (*Actb*) was used as internal control. Boxes represent cropped blots.

To determine if reduction in total cellular O-GlcNAcylation is required for proper erythroid differentiation, we treated GATA-1-ER cells with both E2 and Thiamet G (TMG), a highly selective inhibitor of OGA that blocks the enzymes ability to remove the O-GlcNAc moiety (Yuzwa et al. 2008; Yuzwa et al. 2012; Hastings et al. 2017). Cells treated with TMG lose the ability to remove the O-GlcNAc moiety leading to increased cellular O-GlcNAc levels, and ultimately alters O-GlcNAc cycling. TMG was added at the same time with E2 to initiated differentiation. As expected, western blots show an increase in O-GlcNAc levels following TMG treatment. FOG-1, GATA-1-ER, and GATA-2 protein levels were unaffected by the TMG treatment (**Figure 1b**). Concomitantly, morphologic maturation (**Figure 1c**) and hemoglobin induction (**Figure 1d**) were observed in G1E-ER4 cells treated with E2±TMG. TMG+E2 did not change β^major^-globin gene transcription levels (**Figure 1e**) relative to E2 treatment only. Additionally, we did not observe morphology or (**Figure 1f**) and hemoglobin induction changes in G1E cells treated with E2 ± TMG (**Figure 1g**) indicating these cells did not undergo erythroid differentiation. We conclude that TMG+E2 did not change the cell morphology, hemoglobin induction, or β^major^-globin gene transcription level relative to 30 hrs E2 treatment only.

### GATA-1 Interacts with OGT and OGA in G1E-ER4 cells

Our previous study in K562 cells, an erythroid leukemia cell line, and murine β yeast artificial chromosome chemically inducible dependent bone morrow cells (βYAC CID BMC) demonstrated that OGT and OGA interact with the GATA-1-FOG-1-Mi2β repressor complex at the −566 GATA site of the ^A^γ-globin gene, and that the Mi2β-OGT-OGA interactions change when the γ-globin transcription is altered (Zhang et al. 2016). Next, we assessed whether OGT and OGA interact with GATA-1-ER during E2 induced differentiation by immunoprecipitations experiments (IPs) using antibodies against O-GlcNAc (RL2), OGT, OGA, and GATA-1 in G1E-ER4 cell lysates treated with TMG and/or E2. We found that GATA-1-ER was poorly immunoprecipitated by O-GlcNAc, OGT, and OGA antibodies before E2 treatment. After E2 treatment, O-GlcNAc, OGT, and OGA robustly immunoprecipitated GATA-1-ER (**Figure 2a-c**). O-GlcNAc, OGT and OGA pulled down FOG-1 before and after E2 treatment, and E2 treatment increased the amount of immunoprecipitated FOG-1 with a molecular weight shift suggesting an increase in FOG-1 phosphorylation (Villen et al. 2007; Huttlin et al. 2010) (**Figure 2a-c**). GATA-1 antibody could immunoprecipitate OGT and OGA before and after E2 treatment (**Figure 2d**). These data indicate that the interaction between GATA-1-ER, OGT, OGA, and the O-GlcNAcylation of GATA-1-ER interacting proteins increases during G1E-ER4 cell differentiation.

**Figure 2.**
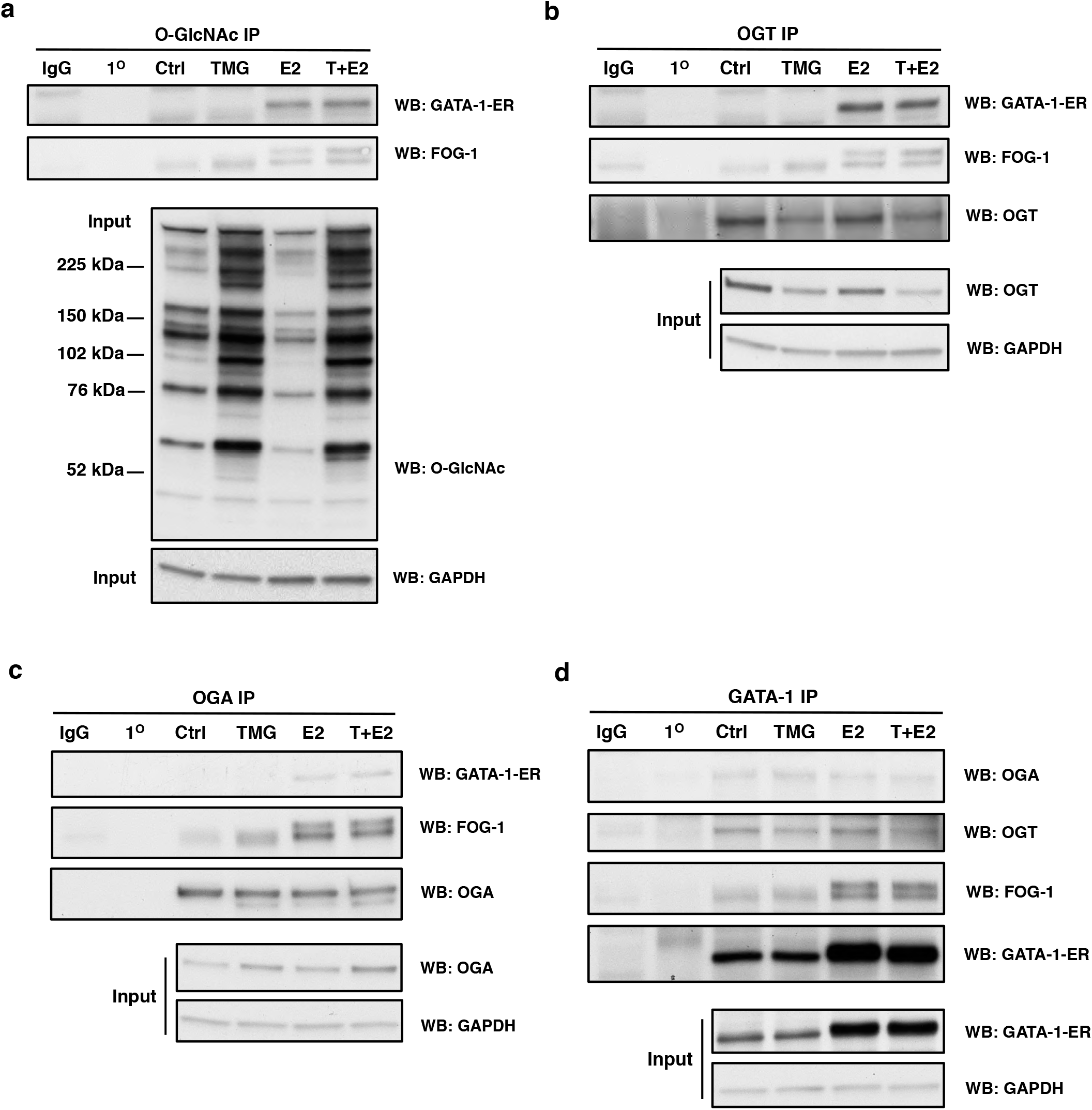
GATA-1 interacts with OGT and OGA in G1E-ER4 cells. G1E-ER4 cells were treated with TMG (T) and/or E2 for 30 hrs. Cells were harvested and subjected to immunoprecipitation (IP). O-GlcNAcylated protein, OGT, OGA, and GATA-1-ER were immunoprecipitated with a specific (**a**) O-GlcNAc antibody (RL2), (**b**) OGT, (**c**) OGA, and (**d**) GATA-1 antibody, respectively. Blots were probed with RL2, GATA-1, FOG-1, OGT, OGA, and GAPDH respectively. Isotype (normal rabbit or mouse IgG) IP and antibody alone precipitation (1°) were used as negative controls. All experiments were performed with at least 3 biological replicates. Boxes represent cropped blots.

### Inhibition of OGA with TMG changes the erythroid gene transcription networks

We have demonstrated that OGT and OGA interact with GATA-1 in multiple erythroid cell lines; therefore, we expanded on these discoveries by testing whether the erythroid gene transcription network was altered by disruptions in O-GlcNAc homeostasis during G1E-ER4 differentiation. We performed Next Generation RNA-sequence analysis on G1E-ER4 cells without treatment, with TMG or E2 treatment only, and E2+TMG treatment for 30 hrs, this time point corresponds with previously published GATA-1 ChIP sequencing data (Cheng et al. 2009). 8,271 transcripts (**Supplemental Table 1**) were included in the RNA-seq analysis from the different treatments, and the shared genes that were down-regulated (**Figure 3a**) or up-regulated (**Figure 3b**) are shown. Our RNA-seq data suggest that inhibition of OGA by TMG changes transcription levels of numerous genes during erythropoiesis. In order to characterize the biological processes altered by inhibition of OGA, Gene Set Enrichment Analysis (GSEA) of the 8,271 transcripts was performed (Mootha et al. 2003; Subramanian et al. 2005). The top 10 up-regulated (top panel) and down-regulated (bottom panel) biological processes are listed comparing TMG+E2 with E2 treatment only (**Figure 3c**). Examples of enrichment analysis plots are shown in **Figure 3d**. Interestingly, biological processes of myeloid leukocyte activation and inflammatory response were up-regulated. To further evaluate our RNA-sequence data, we compared the transcription profile of G1E-ER4 cells treated with E2+TMG to E2 only (**Figure 4a**). 1,173 transcripts (**Supplemental Table 2**) were differentially expressed in E2+TMG compared to E2 treatment only, and 530 genes were up-regulated while 643 were down-regulated (**Figure 4b-c**), including 47 erythroid-specific genes (**Supplemental Table 3**) based on a previous study (Li et al. 2013). These results indicate that many biological processes were changed when OGA was inhibited with TMG during erythropoiesis, and that alteration of O-GlcNAc cycling in GATA-1-ER cells modulates terminal differentiation of erythroid gene programs.

**Figure 3.**
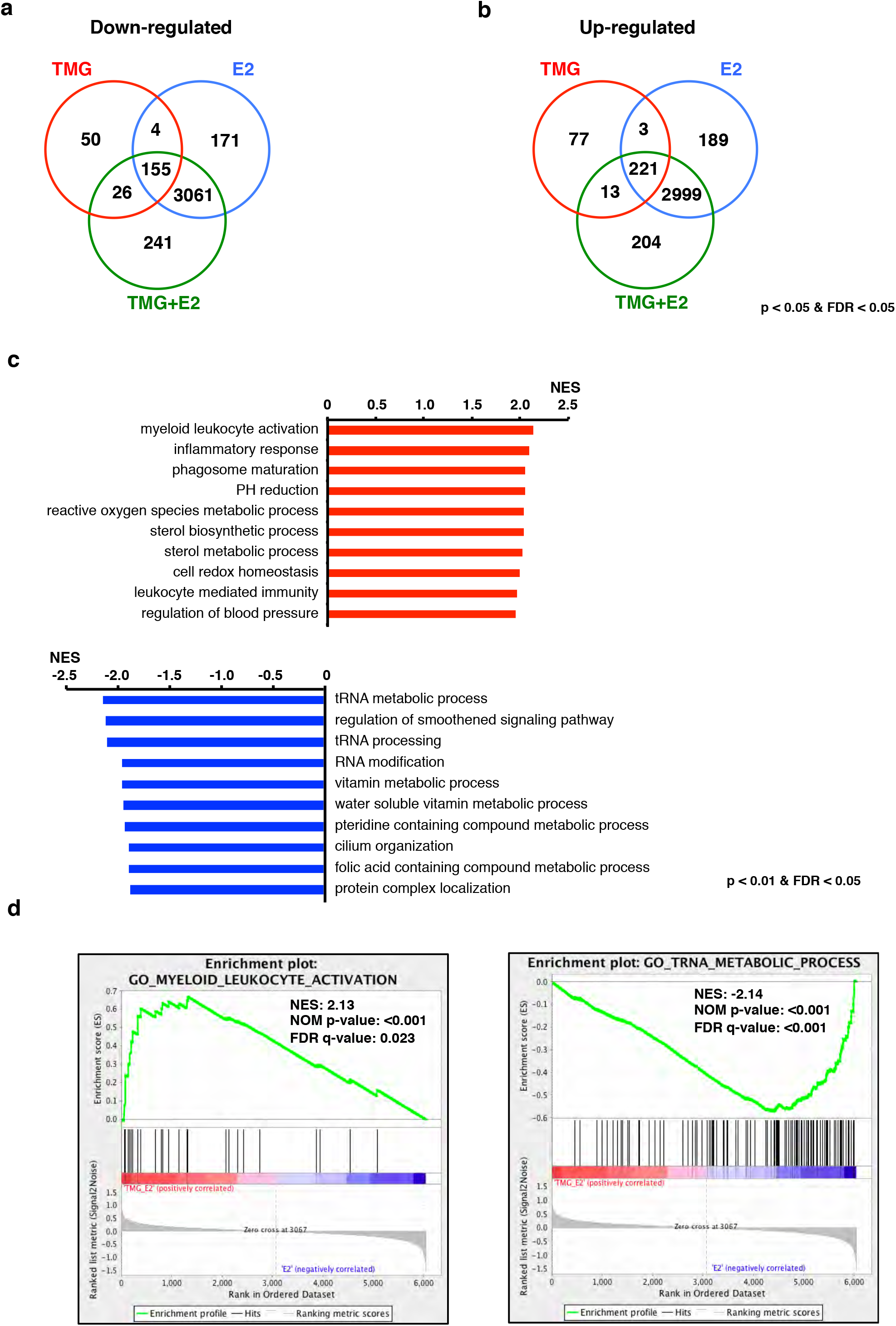
Inhibition of OGA changes gene transcription network during erythropoiesis. Venn diagram of (**a**) down- and (**b**) up-regulated transcripts in TMG, E2, and TMG+E2 treatment compared to control, respectively. Transcripts with p value<0.05 and FDR (False Discover Rate) <0.05 were included in the analysis. **c**. Top 10 up-(top) and down-regulated (bottom) biological processes analyzed by GSEA are ranked by NES (Normalized Enrichment Score). **d**. Enrichment plot of an up-regulated biological process myeloid leukocyte activation (left) and down-regulated biological process tRNA metabolic process (right). NOM (Nominal) p-value < 5% and FDR q-value < 25% were used for biological processes analysis.

### O-GlcNAc cycling modulates GATA-1-ER target gene transcription during erythropoiesis

To evaluate and explore how O-GlcNAc cycling modulates GATA-1-ER target gene transcription, we used an online database, Harmonizome (Rouillard et al. 2016), to search for GATA-1 target genes in G1E-ER4 cells after E2 treatment. A total of 4,072 GATA-1 target genes were selected from the database by combining 9 sets of ENCODE mouse GATA-1 binding site profiles in G1E-ER4 cells (Consortium 2004; Rouillard et al. 2016). 433 of them (**Supplemental Table 4**) are differentially expressed in TMG+E2-treated cells compared to E2 only-treated cells (**Figure 4d**). Of these 433 genes, 243 genes were up-regulated and 190 genes were down-regulated (**Figure 4d**). We again preformed GSEA analysis on these 433 genes. Immune response and defense response biological processes, which are characteristics of leukocytes, were again observed to be up-regulated in TMG+E2-treated cells compared to E2 only-treated cells (**data not shown**).

**Figure 4.**
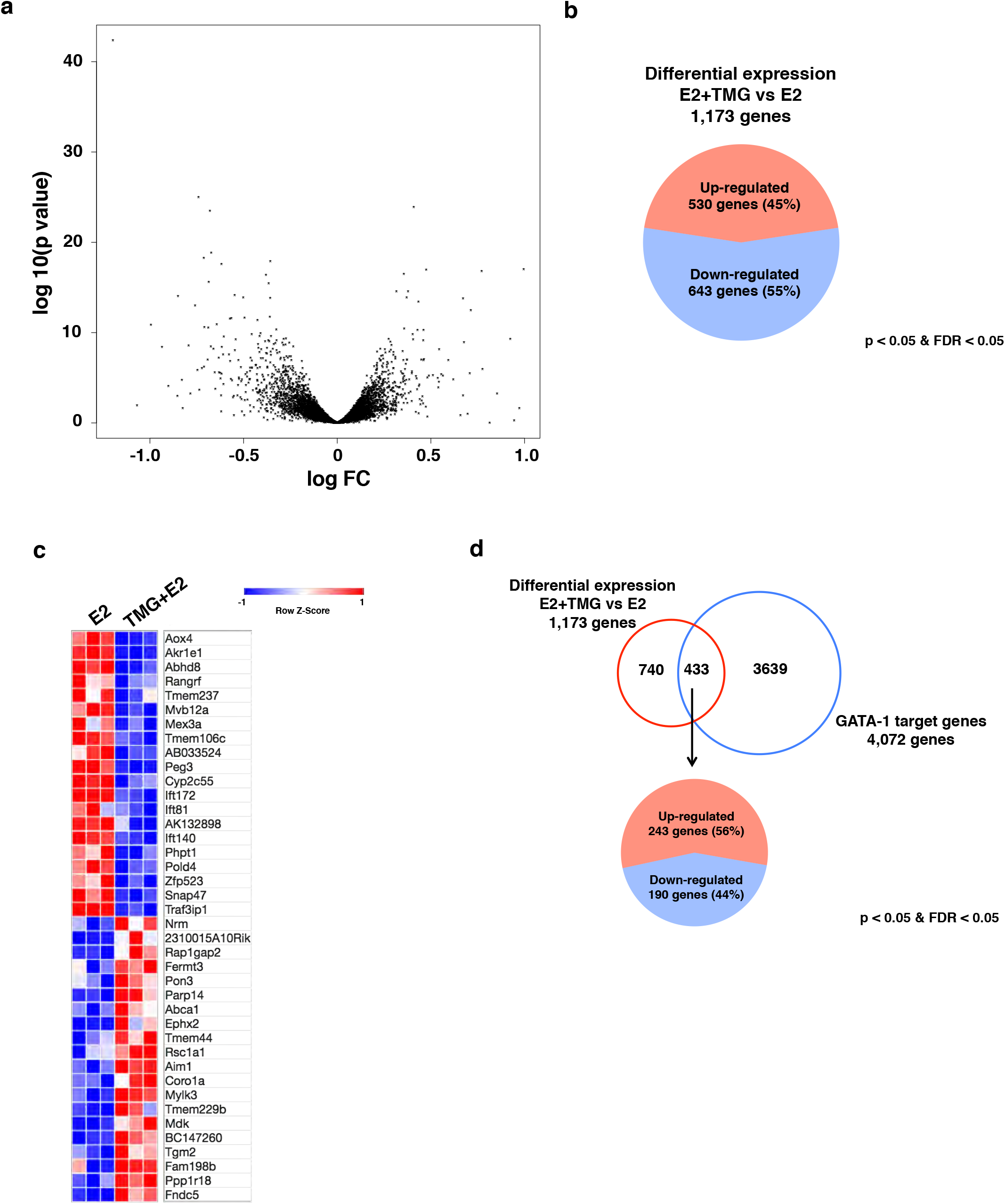
O-GlcNAc regulates GATA-1 target gene transcription during erythropoiesis. **a**. The Volcano Plot of differentially expressed genes in E2+TMG compared to E2 treatment only. Each black dot represents single gene from RNA-seq analysis, with the negative log of P value plotted on the Y-axis and the log of the fold change (FC) plotted on the X-axis. **b**, 1,173 genes were differentially expressed in E2+TMG compared to E2 treatment only. Pie chart shows the distribution of up-(45%) and down-regulated (55%) genes differentially expressed. Transcripts with p value < 0.05 and FDR < 0.05 were included in the analysis. **c**, The heat map of top 20 differentially expressed genes. **d**, Venn diagram of differential expression of 1,173 genes (TMG+E2 versus E2) and 4,072 GATA-1 targeted genes from database. Of 433 differentially expressed GATA-1 target genes, 243 genes were up-regulated and 190 genes down-regulated.

In order to assess the quality of our RNA-seq data, we selected 4 genes for quantitative Polymerase Chain Reaction (qPCR) analysis. *Laptm5, Fndc5, Parp14*, and *Mvb12a* are all GATA-1 target genes showing a robust transcription level change following TMG+E2 treatment compared to E2 only treatment according to our RNA-seq data analysis. Total RNA was extracted from cells under different experimental conditions which included: 1) no treatment (control); 2) TMG or E2 only treatment; and 3) E2+TMG treatment. After E2 treatment for 30 hrs, *Laptm5, Fndc5*, and *Parp14* transcription levels were increased compared to control (**Figure 5a-c**), while *Mvb12a* transcription level decreased (**Figure 5d**). When cells were treated with TMG+E2, *Laptm5, Fndc5*, and *Parp14* transcription levels were increased compared to E2 only treatment (**Figure 5a-c**), while *Mvb12a* transcription decreased (**Figure 5d**). These results were consistent with the fold changes from RNA-seq data. We repeated identical measurements on expression of these genes in G1E cells (negative control); no change was detected in expression with TMG, E2, or E2+TMG (data not shown). This data confirms that a subset of GATA-1 target genes was affected by alterations in O-GlcNAc homeostasis during erythropoiesis.

**Figure 5.**
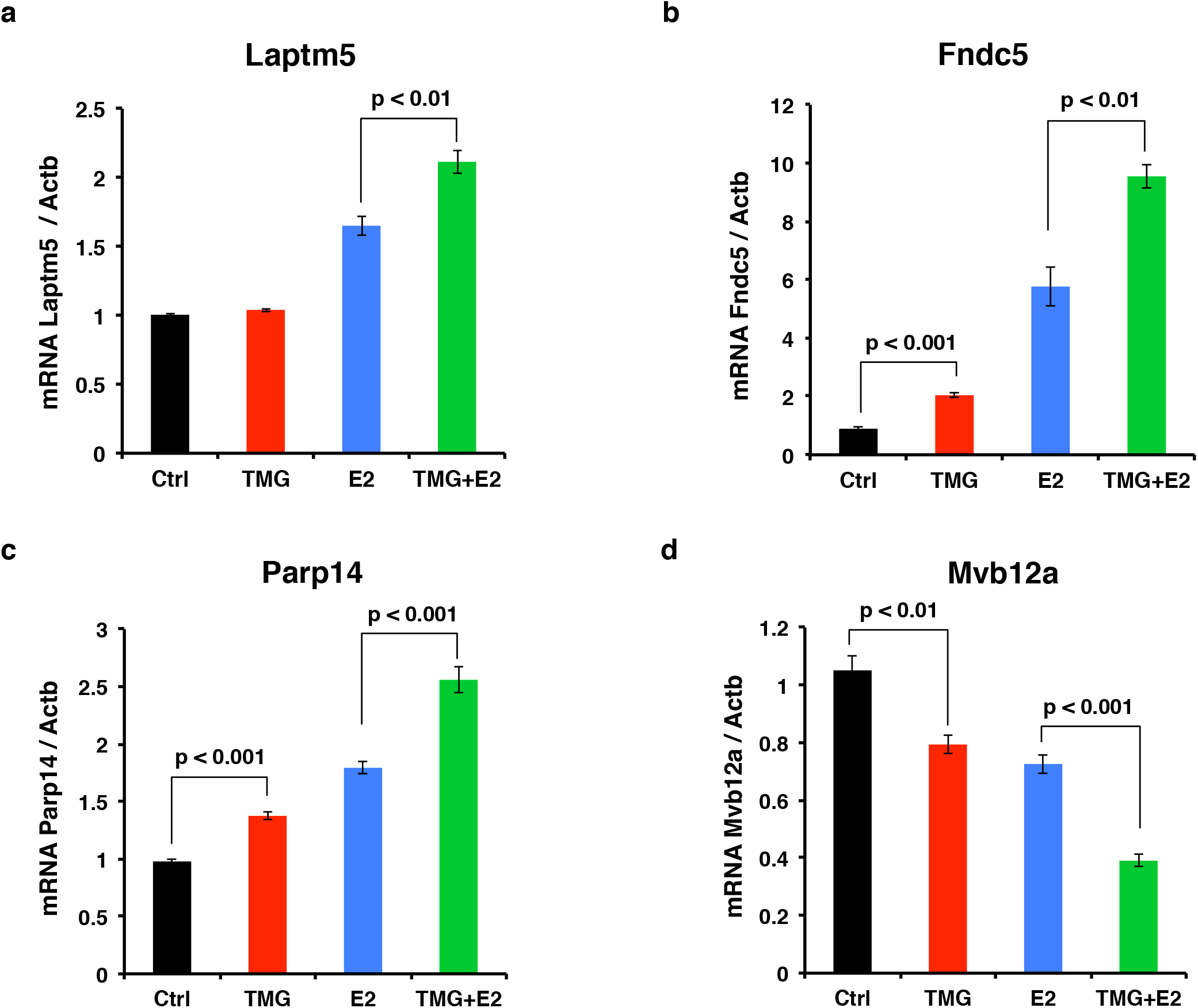
Validation of O-GlcNAc regulated GATA-1 target gene by qPCR. Transcription level of GATA-1 target genes (**a**) *Laptm5*, (**b**) *Fndc5*, (**c**) *Parp14*, and (**d**) *Mvb12a* was analyzed by qPCR after cells were treated with TMG and/or E2. The experiments were repeated 4 times. β-Actin (*Actb*) was used as internal control.

### Inhibition of OGA decreases the occupancy of GATA-1, OGT, OGA, and O-GlcNAc levels at the *Laptm5* GATA binding site

Based on our RNA-seq and GSEA analysis, we observed that inhibition of OGA by TMG altered a subset of GATA-1 target genes during erythropoiesis, in particular immune response and defense response genetic pathways. *Laptm5*, a target of GATA-1-ER activation, is specifically expressed in hematopoietic cells, and is preferentially expressed in immune cells (Adra et al. 1996; Glowacka et al. 2012a). LAPTM5 positively modulates inflammatory signaling pathways and cytokine secretion in macrophages (Glowacka et al. 2012b), and is up-regulated in G1E-ER4 after 30 hrs E2 treatment. We performed ChIP assays at the GATA binding site located in the first intron of *Laptm5*, approximately 8 kb downstream of the transcription start site (TSS) (Cheng et al. 2009) using antibodies against GATA-1, OGT, OGA, and O-GlcNAc to assess the occupancy of GATA-1-ER, OGT, OGA, and O-GlcNAc. In untreated control G1E-ER4 cells, we observed the occupancy of OGT, OGA, and an enrichment of O-GlcNAcylation at the +8 kb *Laptm5* GATA binding site (**Figure 6b-d**), suggesting that O-GlcNAcylation at the *Laptm5* GATA binding site might be important for maintaining the basal level of *Laptm5* transcription. Following GATA-1-ER restoration by E2, GATA-1-ER occupancy dramatically increased at the GATA binding site compared to the untreated control; however, E2+TMG reduces the amount of GATA-1-ER occupancy at this site (**Figure 6a, left panel**). OGT occupancy did not change and OGA occupancy increased at the GATA binding site after E2 treatment compared to control (**Figure 6b and c, left panel**). We hypothesize that this contributes to the decrease in overall O-GlcNAc levels at this GATA binding site (**Figure 6d, left panel**). Interestingly, the occupancy of OGT and OGA, and the overall O-GlcNAc level at this GATA binding site decreased after TMG+E2 treatment compared to E2 only treatment (**Figure 6b-d, left panel**). A similar pattern was observed when TMG only treatment was compared with the untreated control (**Figure 6b-d, left panel**). To show that these changes were specific to the GATA binding site, we used a non-GATA-1 binding DNA region 50 kb downstream of the GATA binding site as a negative control. We did not observe any changes when comparing TMG+E2 treatment with E2 only treatment (**Figure 6a-d, right panel**). This data suggest that O-GlcNAc plays a role in GATA-1 regulation of the *Laptm5* gene and that TMG treatment alters the occupancy of GATA-1-ER, OGT, and OGA at this binding site.

**Figure 6.**
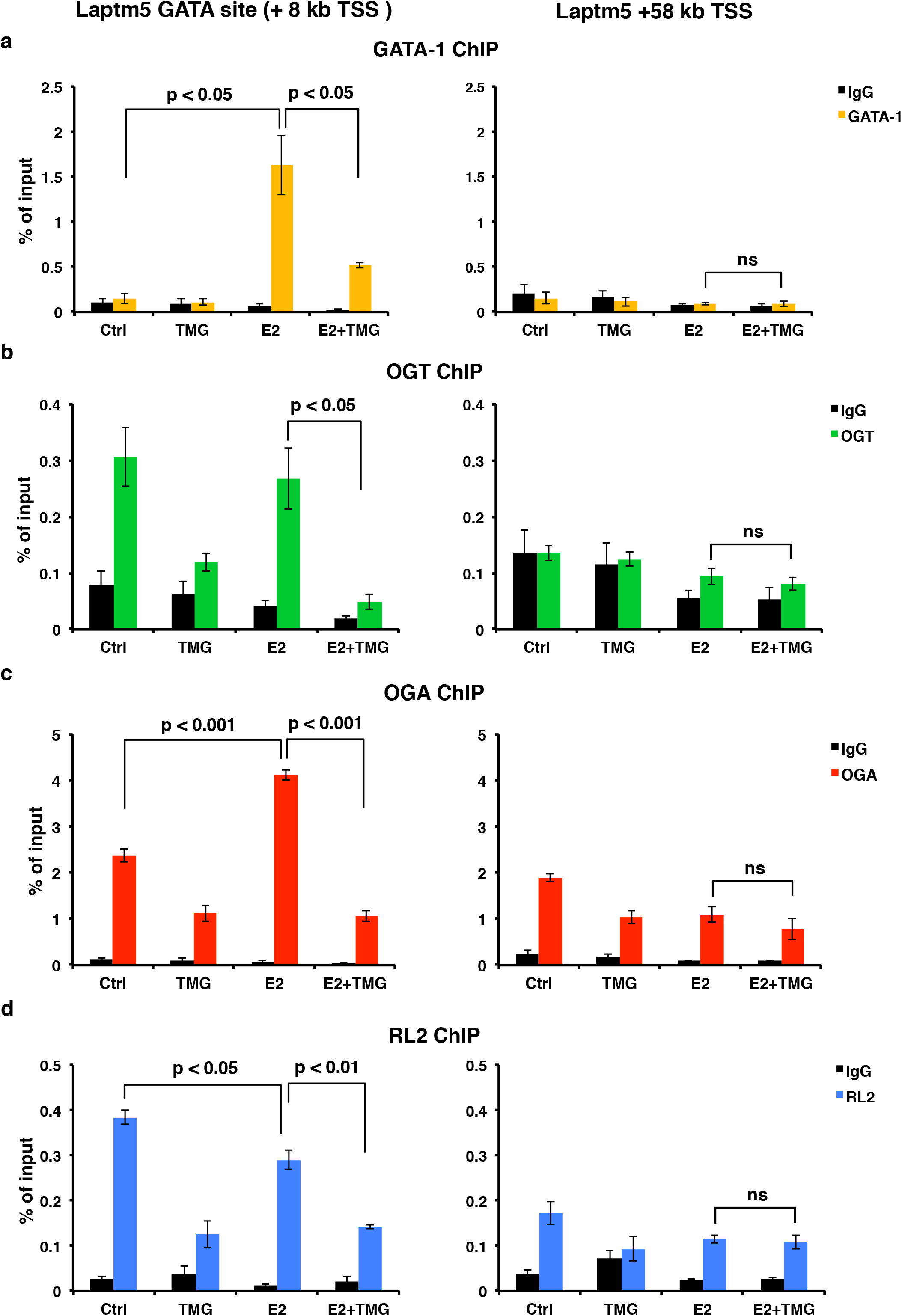
Inhibition of OGA decreases the occupancy of GATA-1-ER, OGT, OGA, and the O-GlcNAc level at *Laptm5* GATA binding site. (**a**) GATA-1, (**b**) OGT, (**c**) OGA, and (**d**) O-GlcNAc (RL2) ChIP assays were performed using G1E-ER4 cells treated with TMG and/or E2. ChIP DnA was analyzed by qPCR using a set of primer targeting the *Laptm5* GATA binding site (+ 8 kb tSs) and +58 kb TSS respectively. Normal rabbit IgG served as a negative control. All experiments were performed with at least 3 biological replicates. ns, not significant.

### Neutrophil-like differentiation is impaired with TMG treatment

Next, we hypothesized that O-GlcNAcylation is involved in the differentiation of other hematopoietic cell lines. NB4 and HL60 myeloid leukemia cells differentiate into neutrophil like cells with the addition of all-trans retinoic acid (ATRA) (Drach et al. 1993). Therefore, we treated cells with ATRA for 48 hrs prior to harvesting. In both cell lines, we measured a decline in O-GlcNAc levels at 48 hrs. The O-GlcNAcylation change in the NB4 cells affected a large number of O-GlcNAc modified proteins, while in HL60 cells proteins at 76, 44, 37, and 12 kDa showed declines in O-GlcNAc (**Figure 7a**). As with the G1E-ER4 cells, OGT levels did not decline with differentiation suggesting that OGT substrate targeting proteins appear to be changing with ATRA induction. Next, we treated NB4 cells with TMG at the time of induction with ATRA and measured the mRNA expression of Cathepsin D (*CTSD*) and Defensin Alpha 1 (*DEFA1*) genes, which increase with neutrophil-like differentiation (Baines et al. 2011; Yoon et al. 2013). ATRA treatment led to a robust increase in *CTSD* (**Figure 7b**) and *DEFA1* (**Figure 7c**) mRNA expression. TMG treatment significantly decreased *CTSD* and *DEFA1* gene expression after ATRA addition. These data phenocopy gene expression changes during G1E-ER4 cells differentiation and suggest that O-GlcNAc levels decline as hematopoietic cells differentiate, substrate targeting of OGT and OGA changes during differentiation, and that disruption in O-GlcNAc homeostasis impairs differentiation.

**Figure 7.**
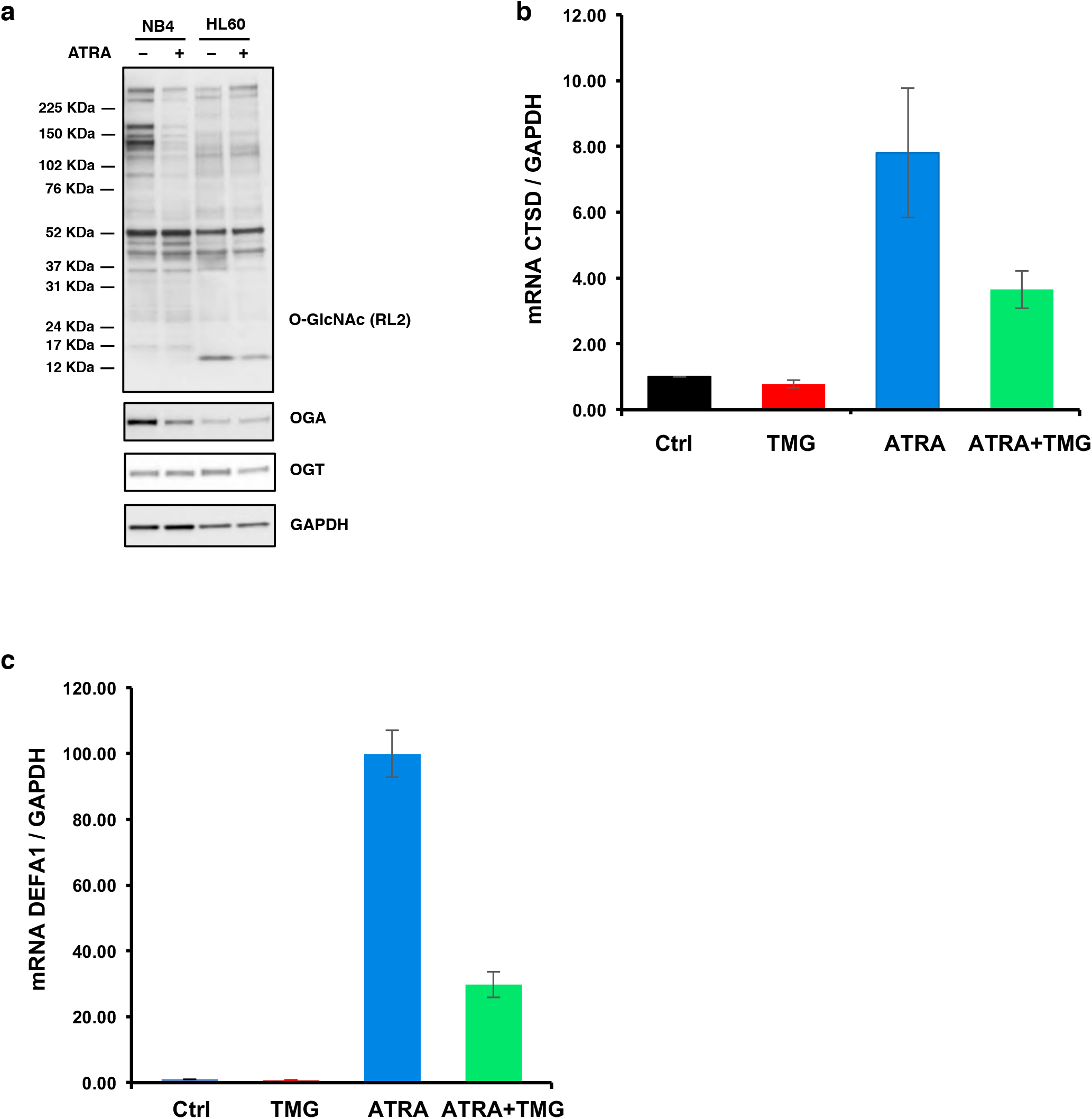
Neutrophil-like differentiation is altered with TMG treatment. NB40 or HL60 cells were induced to differentiate with ATRA + TMG. Cells were harvested at 48 hrs and total cell lysates were subjected to immunoblotting (**a**). Transcription level of genes induced by ATRA treatment (**b**) *CTSD* and (**c**) *DEFA1* were analyzed by qPCR. The experiments were repeated 3 times. β-Actin (*ACTB*) was used as internal control.

### Prolonged OGA inhibition impairs erythropoiesis

Previously, we demonstrated in both cell lines and mice tissue that sustained TMG treatment (14 days or longer) alters the metabolic demands of a cell and reprograms the transcriptome leading to a variety of adaptive changes to TMG (Tan et al. 2017). Sustained TMG alters O-GlcNAc homeostasis by elevating O-GlcNAc to a new set point and increasing OGA expression to compensate (Zhang et al. 2014).

Therefore, we treated G1E-ER4 cells for 2 weeks with a daily dose of TMG and then differentiated the cells with E2. At 30 hrs post-differentiation, there was no difference between E2, E2+TMG, or E2 with sustained TMG treatment in total hemoglobin positive cells or % of cells alive. At 76 hrs post-E2 treatment, E2 treatment only and acute TMG+E2 treatment has similar number of cells that were hemoglobin positive and alive; however sustained TMG-treated cells had a far higher proportion of cells that were alive and far less cells positive for hemoglobin suggesting that the alteration in O-GlcNAc homeostasis induced by the adaption to TMG was slowing differentiation (**Table 1**). Next, we measured β-globin expression at 76 hrs post-differentiation with E2 and sustained TMG treatment. β^major^-globin expression was dramatically increased at 76 hrs after E2 treatment compared to control, but sustained TMG+E2 treatment reduced β^major^-globin expression by half compared to E2 treatment only (**Figure 8a**). However, the *Fndc* gene was increased with sustained TMG treatment and E2 (**Figure 8b**) and *Laptm5* had no changes in all groups at 76 hrs (**data not shown**). These data demonstrate that TMG treatment slows G1E-ER4 differentiation and alters the expression of erythroid genes.

**Figure 8.**
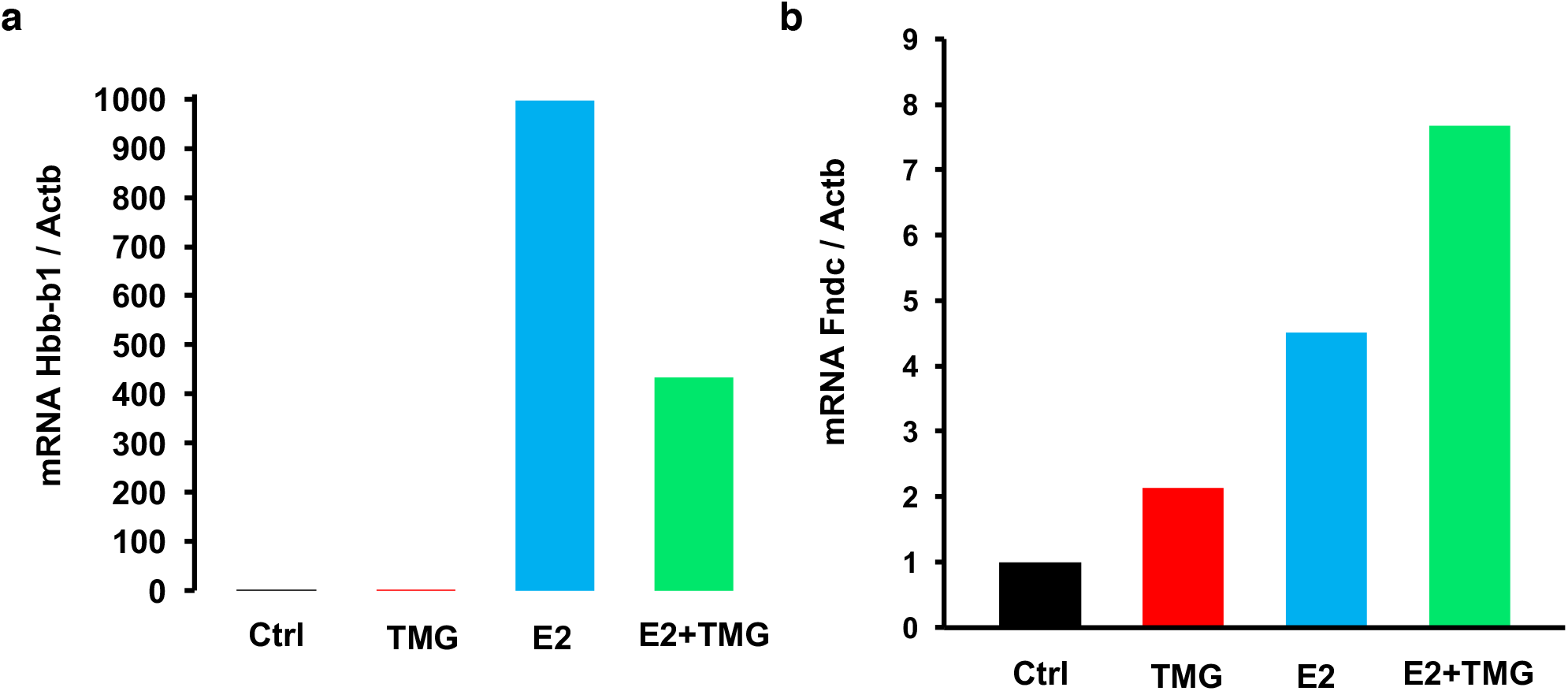
Sustained OGA inhibition modulates erythropoietic gene expession. G1E-ER4 cells were treated with TMG (for 2 weeks) and/or E2 for 76 hrs. Cells were harvested. Transcription level of (**a**) β^major^-globin gene (*Hbb-b1*) and (**b**) *Fndc5* was measured by qPCR and β-actin (*Actb*) was used as internal control.

**Table 1.** Total % of hemoglobin containing live cells with E2, acute, and long term TMG treatment

| TMG treatment<br>Induction time | no TMG treatment + E |  |  | acute TMG treatment + E |  |  | 2 weeks TMG treatment + E |  |  |
| --- | --- | --- | --- | --- | --- | --- | --- | --- | --- |
| Estradiol, hours | Live, blue, % of total | Hb positive, brown, % of total | Hb positive, gray-blue, % of total | Live, blue, % of total | Hb positive, brown, % of total | Hb positive, gray-blue, % of total | Live, blue, % of total | Hb positive, brown, % of total | Hb positive, gray-blue, % of total |
| 30 | 91.17 | 3.59 | 5.24 | 90.90 | 6.17 | 2.93 | 91.75 | 5.32 | 2.93 |
| 76 | 45.97 | 42.90 | 11.13 | 47.81 | 42.36 | 9.83 | 73.39 | 19.64 | 6.96 |

## Discussion

In this study, we took advantage of a well-established cell model of erythropoiesis, G1E-ER4 cells, to investigate the role of O-GlcNAc during GATA-1-mediated erythroid differentiation. After restoration of GATA-1 activity, overall O-GlcNAc levels dramatically decreased, with an increase of OGA and a decrease of OGT protein levels. The decrease in O-GlcNAcylation during differentiation was also seen in two myeloid leukemia cell lines when they were induced to become more neutrophil-like. In addition, GATA-1-ER restoration promoted the interaction of GATA-1-ER with OGT and OGA. RNA-seq analysis of G1E-ER4 cells treated with E2 and TMG demonstrated a change in transcription level of 1,173 genes compared to cells treated with E2 only, including 433 GATA-1 target genes. The transcription level changes were orthogonally validated on selected genes by qPCR. One of these genes *Laptm5* was subjected to further proof-of-principle analysis. ChIP data demonstrated that the occupancy of GATA-1 and oGa was increased and overall O-GlcNAc level was decreased at the *Laptm5* GATA binding site following GATA-1 activation, and that these changes were diminished by inhibition of OGA with TMG. TMG treatment impaired expression of neutrophil genes during ATRA mediated differentiation of myeloid cells. Interestingly, sustained TMG treatment in G1E-ER4 reduced β^major^-globin expression but increased expression of *Fndc5*. Together, these data suggest that O-GlcNAc plays a role in regulating a subset of GATA-1 controlled erythroid genes and disruptions in O-GlcNAc homeostasis impairs hematopoietic cell differentiation.

We previously demonstrated that GATA-1 interacts with OGT and OGA in MEL BirA cells (Zhang et al. 2016), and our current results validate these findings. We show that in G1E-ER4 cells, more GATA-1-ER was co-immunoprecipitated after E2 treatment by O-GlcNAc, OGT, or OGA antibodies, respectively, compared to cells without E2 treatment. We did not find that GATA-1 itself was modified by O-GlcNAc. However, GATA-1 interacting proteins are likely modified by O-GlcNAc, since following differentiation, the O-GlcNAc antibody pulled down GATA-1-ER despite the decrease of overall O-GlcNAc level. In our previous study, we also demonstrated GATA-1 interacting protein Mi2β is O-GlcNAc-modified (Zhang et al. 2016). The alteration of OGT and OGA interaction with GATA-1-ER and O-GlcNAcylation of GATA-1-ER interacting proteins during erythropoiesis could change protein-protein interactions within GATA-1-containing activator or repressor complexes, thereby contributing to GATA-1-mediated gene activation or repression. Regulation of transcription factors and co-activator/co-repressor complexes by O-GlcNAc is poorly understood. OGT is part of the mSin3A complex (Yang et al. 2002) and the polycomb complex (Chu et al. 2014; Maury et al. 2015); and both OGA and OGT help organize the RNA Pol II pre-initiation complex (Ranuncolo et al. 2012; Lewis et al. 2016; Resto et al. 2016). Recent advances in O-GlcNAc crosslinking technology will aid in the determination of which proteins interact with O-GlcNAcylated transcriptional complexes, how these complexes change in response to different stimuli, and potentially identify protein domains that preferentially bind to O-GlcNAcylated moieties (Rodriguez et al. 2015).

Our RNA-seq analysis suggests that inhibition of OGA by TMG alters gene transcription during erythropoiesis, and that these changes may reflect reprogramming of the cell to a more myeloid leukocyte-like cell morphology or behavior. Thus, it would be interesting to explore whether O-GlcNAc plays a role in determining hematopoietic cell fate at an earlier stage. For example, does inhibition of OGA by TMG in hematopoietic stem cells (HSCs) prior to lineage commitment change the O-GlcNAcylation pattern or skew HSC differentiation towards a particular lineage? Previous reports show that elevated O-GlcNAc in mouse embryonic stem cells delays the onset of neuronal differentiation (Speakman et al. 2014), while increased O-GlcNAcylation skews human pluripotent stem cell differentiation toward adipose mesoderm markers and away from ectoderm markers (Maury et al. 2015). Since myeloid leukemia cells also show reduced O-GlcNAcylation during differentiation, then OGA inhibition might reduce hematopoietic differentiation in general, or shift commitment towards another hematopoietic lineage. Differentiating primary human CD34+ cells in the presence of TMG might elucidate whether OGA inhibition drives hematopoietic stem cells toward a specific linage and we will explore this hypothesis in future studies.

Of the 1,173 differentially expressed genes (TMG+E2 compared to E2 only), we determined that 47 of these genes are erythroid-specific genes, including *Gfi1b* (Saleque et al. 2002), *Alas2* (Nakajima et al. 1999), and *Gata2* (Orkin 1992), which are critical for normal erythroid differentiation. Crucially *Gfi1b* is a transcription factor important in promoting erythrocyte differentiation; *Gata2* inhibits erythrocyte differentiation, while *Alas2* is critical in porphyrin biosynthesis. We also found 433 of the differentially expressed genes were under control of GATA-1, with 243 genes up-regulated and 190 down-regulated. Notably, 84% of activated GATA-1 target genes became more activated in the presence of tMG; while 79% of repressed GATA-1 target genes became more repressed in the presence of TMG indicating that inhibition of OGA amplifies GATA-1 mediated transcriptional effects. Recently, we showed that in immortalized murine bone marrow cells carrying the human β-globin locus on a yeast artificial chromosome (β-YAC), transcription of the ^A^γ-globin gene was repressed by a GATA-1-FOG-1-Mi2β repressor complex, and that TMG treatment of these cells enhanced the repression of ^A^γ-globin transcription (Zhang et al. 2016).

One of the known GATA-1 target genes is *Laptm5*, which is expressed in hematopoietic cells (Adra et al. 1996). LAPTM5 positively modulates inflammatory signaling pathways and cytokine secretion in macrophages (Glowacka et al. 2012b). The transcription of *Laptm5* was activated upon GATA-1 restoration in G1E-ER4 cells and the activation was enhanced in the presence of TMG. Strikingly, the occupancy of GATA-1-ER, OGT, and OGA, and the O-GlcNAc level at the *Laptm5* GATA binding site decreased after inhibition of OGA. Even though GATA-1-ER occupancy at the *Laptm5* GATA binding site was decreased following TMG+E2 treatment, TMG amplified the increase of *Laptm5* transcription compared to E2 treatment only. Thus, TMG treatment and subsequent retention of O-GlcNAc-modified proteins may stabilize the interaction of GATA-1 with other transcription factors, thereby reducing the recruitment or turnover of these transcription factors. Hence, even with reduced occupancy by GATA-1 at the GATA binding site, the GATA-1 complex present may be more stable or more active in promoting transcription. These data further argue for the importance of the rate of O-GlcNAc cycling at GATA binding sites. Our results mirror RNA Pol II PIC formation. O-GlcNAcylation of RNA Pol II C-terminal domain (CTD) by OGT promotes PIC formation but gene transcription cannot continue until OGA is recruited to the promoter to remove O-GlcNAc on the RNA Pol II CTD, which in turn is activated by phosphorylation (Ranuncolo et al. 2012). The reduced O-GlcNAc cycling at the *Laptm5* gene with TMG+E2 treatment may lead to the recruitment of other transcriptional co-activators or stabilize the GATA-1-containing activator complex at the *Laptm5* GATA binding site resulting in increased transcription.

Based on our data, we propose the following model for OGT, OGA, and GATA-1 regulation of *Laptm5* transcription during erythropoiesis. Following GATA-1 restoration by E2, a GATA-1 transcriptional activator complex is recruited to the GATA site located in the first intron of *Laptm5* (Consortium 2004; Cheng et al. 2009), resulting in increased *Laptm5* gene expression (**Figure 9a**). We propose that OGT interacts with and modifies a component(s) of this complex, thereby promoting the recruitment of other transcriptional co-activators. OGA then removes O-GlcNAc from one or more of the proteins in the complex to stabilize the complex. When OGA is inhibited by TMG, O-GlcNAc cycling at the activator complex is disrupted, leading to decreased occupancy of OGT, OGA, and GATA-1 at the GATA binding site. In addition, it is possible that the components of the activator complex might also change in the absence of OGA function (**Figure 9b**). Regardless, we hypothesize that this alternate activator complex is likely to be more stable and active due to the changes in protein-protein interactions within the complex or alterations of the nearby chromatin structure, resulting in enhanced transcription of *Laptm5* (**Figure 9c**). These data suggest that inhibition of OGA by TMG might affect transcription of a subset of GATA-1 target genes by enhancing gene activation or repression via a mechanism yet to be determined.

**Figure 9.**
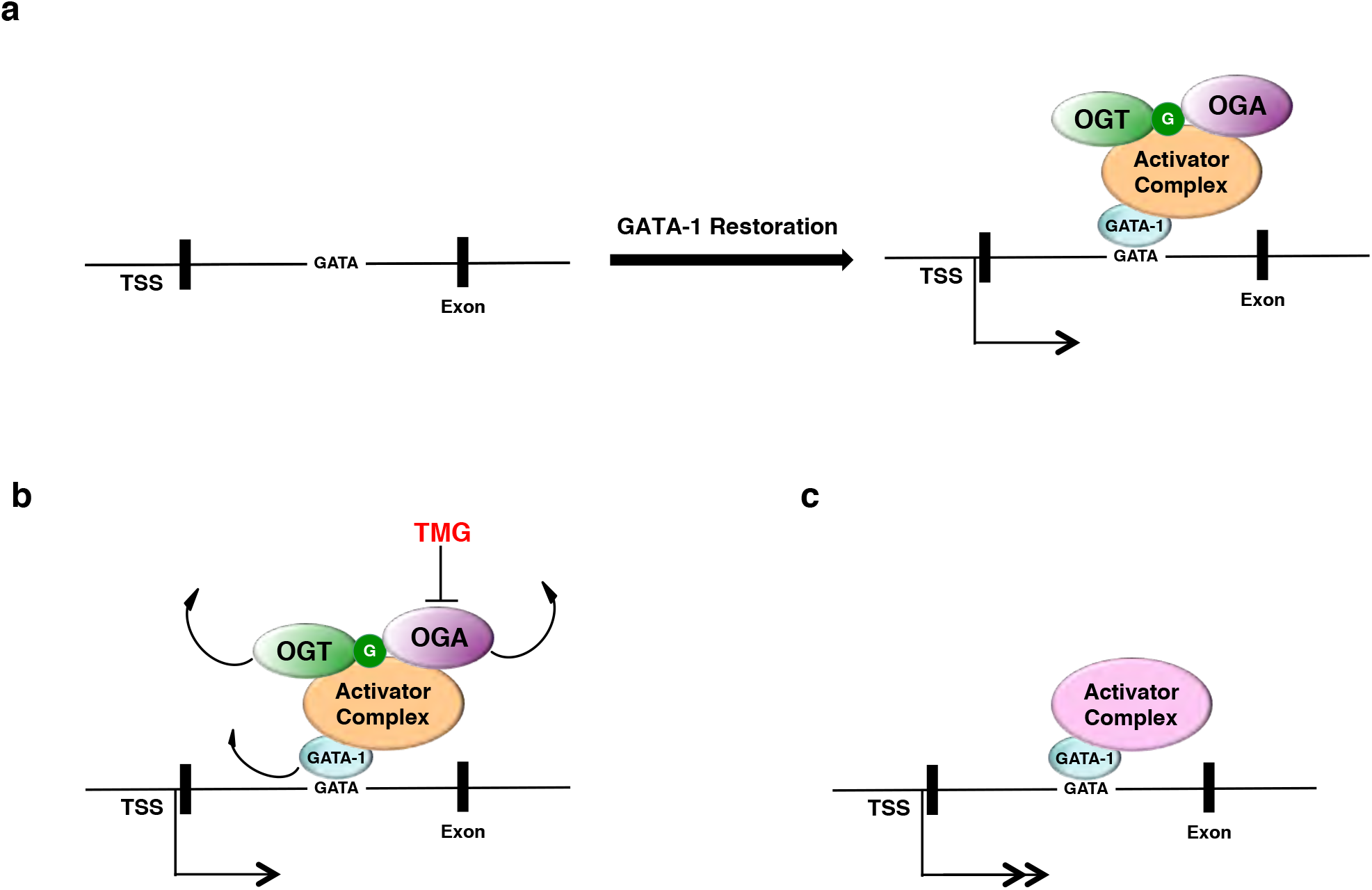
Proposed mechanism of OGT and OGA mediated repression and activation of GATA-1 target gene *Laptm5* during erythropoiesis. In untreated G1E-ER4 cells, *Laptm5* transcription is maintained at a basal level. Following GATA-1 restoration by E2, a GATA-1 transcriptional activator complex (orange oval) is recruited to the GATA site located in the first intron of *Laptm5*, resulting in increased *Laptm5* gene expression (**a**). When OGA is inhibited by TMG, O-GlcNAc cycling at the activator complex (orange oval) is disrupted, leading to decreased occupancy of OGT, OGA, and GATA-1 at the GATA binding site. In addition, it is possible that the components of the activator complex might also change in the absence of OGA function (**b**). This alternate activator complex (pink oval) is likely to be more stable and active due to the changes in protein-protein interactions within the complex or alterations of the nearby chromatin structure, resulting in enhanced transcription of *Laptm5* (**c**).

Based on our data we hypothesize, O-GlcNAcylation is required for hematopoietic cell differentiation. Our data does recapitulate data from other studies of differentiation (Kim et al. 2009; Ogawa et al. 2012; Sohn et al. 2014; Speakman et al. 2014; Zhang et al. 2016), which also show a decrease of overall O-GlcNAcylation, suggesting that in most cells, differentiation requires a reduction of O-GlcNAc levels. The relationship of reduced O-GlcNAc levels to differentiation deserves further study. Many cells enter the cell cycle prior to differentiation suggesting that a decrease in O-GlcNAc might enhance cell cycle progression (Slawson et al. 2005). Alternately, cells might redistribute metabolites away from the hexosamine biosynthetic pathway which makes UDP-GlcNAc, the metabolic substrate for OGT, to other metabolic pathways needed for growth (Ferrer et al. 2016). We posit that O-GlcNAcylation is a key regulator of transcriptional programs involved with linage specific differentiation. Since O-GlcNAcylation regulates transcription factor, RNA Polymerase II, and chromatin remodeler function in addition to being an epigenetic mark (Hardiville and Hart 2014), then alterations to O-GlcNAc homeostasis would have a dramatic effect on gene programs. Crucially, disruptions in O-GlcNAc homeostasis caused by disease or nutrient excess would stress hematopoietic cell differentiation.

## Materials and Methods

### Antibodies

Primary antibodies and secondary antibodies for immunoblotting were used at 0.5 μg/ml and 1:10,000 dilution respectively. Antibodies for immunoprecipitation (IP) and chromatin immunoprecipitation (ChIP) assay were used at 2 μg per reaction. Anti-OGT (AL-34) and OGA (345) were gracious gifts from Dr. Gerald Hart at the Johns Hopkins University School of Medicine. Anti-O-linked N-acetylglucosamine antibody (RL2) (12440061), and rabbit anti-goat IgG HRP (31402) were purchased from ThermoFisher Scientific. Anti-GAPDH antibody (ab9484) and anti-GATA2 antibody (ab109241) were purchased from Abcam. Anti-chicken IgY HRP (A9046) was purchased from Sigma. Goat anti-rabbit IgG HRP (170-6515) and goat anti-mouse IgG HRP (170-6516) were purchased from Bio-Rad. Goat anti-rat IgG HRP (NA935V) was purchased from GE Healthcare. GATA-1 antibody (sc-265 X) and FOG antibody (sc-9361 X) were purchased from Santa Cruz Biotechnology. Antibodies exclusive for IP: Normal rabbit IgG (sc-2027), normal mouse IgG (sc-2025), and rat IgG (sc-2026) were purchased from Santa Cruz Biotechnology. Anti-OGT (dM-17) antibody (O6264) and anti-OGA antibody (SAB4200267) were purchased from Sigma. Antibodies for exclusive for ChIP: Rabbit control IgG (ab46540) and anti-GATA-1 antibody (ab11852) were purchased from Abcam. ChIP grade mouse (G3A1) mAb IgG1 isotype control (5415) was purchased from Cell Signaling Technologies. OGT antibody (61355) was purchased from Active Motif.

### Cell Culture

G1E (<u>G</u>ATA-<u>1</u>^−^ <u>E</u>rythroid) and G1E-ER4 cells were kind gifts from Dr. Soumen Paul at the University of Kansas Medical Center. These lines were cultured as described previously (Welch et al. 2004; Grass et al. 2006). Cells were incubated at 37°C, 5% CO_2_ in a 95% humidified incubator. Cells were treated with 10 μM Thiamet-G (TMG, S.D. Specialty Chemicals) and/or 1 μM β-estradiol (E2, E2758, Sigma) (Grass et al. 2006). After 30 hrs, cells were harvested for subsequent analysis. For sustained TMG treatment, TMG was added to cells everyday (10 μM) for 2 weeks prior to E2 induction. NB4 ad HL60 cells were grown as previously described (Wang et al. 2011). ATRA (R2625, Sigma) was used at 1 μM in DMSO.

### Western Blot and Immunoprecipitation (IP)

Cells were lysed on ice as described previously (Zhang et al. 2016). Total cell lysates were used for electrophoresis and subjected to Western blot as previously described (Zhang et al. 2016). All Western blot results were repeated in three independent experiments and representative images are shown. IP was performed as described previously (Zhang et al. 2016). Two mg of cell lysates and 2 μg of antibody were used in each reaction.

### Benzidine Staining and May-Grünwald/Giemsa Stain

Benzidine Staining and May-Grünwald/Giemsa Stain were performed as described previously (Welch et al. 2004) to assess the hemoglobin content and morphology changes, respectively. Cell stain images were visualized and photographed using a Nikon Eclipse 80i digital microscope (Nikon Instruments INC, Melville, NY).

### Total RNA Isolation and RT-PCR

Total RNA was isolated from 5 × 10^6^ cells using TRI Reagent (T9424, Sigma) according to the manufacturer’s instructions. For reverse transcription (RT), 0.5 μg of total RNA was reverse-transcribed to cDNA using iScript Reverse Transcription Supermix (170-8841, Bio-Rad) following the manufacturer’s instructions. Reactions were incubated in a thermal cycler (Model 2720, Applied Biosystems) using the following protocol: priming 5 min at 25°C, RT 20 min at 46°C, RT inactivation 1 min at 95°C, and hold at 4°C.

### RNA-Sequencing

RNA was extracted using the Tri Reagent and RNA sequencing libraries were prepared using the Illumina TruSeq Stranded mRNA Library Preparation Kit Set A (RS-122-2101, Illumina) according to manufacturer’s instructions. RNA libraries were sequenced using an Illumina HiSeq 2500 Sequencing System.

### RNA-sequencing Data Analysis

FastQC (0.11.2) and RSEM (1.2.22) software were used to assess the quality of the RNA sequencing results, align the reads to the mouse genome reference GRCm38/mm10, and calculate the gene expression values. R (3.2.2) and EdgeR (3.14.0) software were used to normalize the expression values using the TMM-method (weighted trimmed mean of M-values), followed by differential expression analysis. To reduce the burden of multiple testing in differential gene expression analyses, a filter was initially applied to reduce the number of genes. Genes were removed if they did not present a meaningful gene expression across all samples; only genes with cpm (counts per million) of >10 in least two samples were considered in differential expression analyses. The Benjamini and Hochberg procedure was used to control the false discovery rate (FDR). The following R-packages were utilized for calculations and visualizations: edgeR and Gplot.

### Chromatin Immunoprecipitation (ChIP) Assay

The ChIP assay was performed using a previously published method (Nelson et al. 2006; Zhang et al. 2016). Briefly, chromatin was prepared and sheared as described (Zhang et al. 2016). For each reaction, sheared chromatin was diluted with IP buffer (150 mM NaCl, 5 mM EDTA, 0.5% NP-40, 1% Triton X-100, 40 mM GlcNAc, and 50 mM Tris-HCl, pH 7.5), supplemented with protease inhibitor, and incubated with 2 μg of control IgG or specific antibody, respectively, at 4°C overnight. Dynabeads Protein G (10004D, ThermoFisher Scientific) were added to the mixture, followed by rotation at 4°C for 2 hrs. Dynabeads were washed with 1 ml cold IP buffer 6 times, then mixed with 10% Chelex 100 slurry (1421253, Bio-rad); for input samples, 10% of sheared chromatin was used and processed at the same time as the ChIP samples. Samples were boiled at 95°C for 10 min, followed by treatment with RNase A and proteinase K. Samples were boiled again for 10 min to inactivate proteinase K and centrifuged at 12,000xg for 1 min at 4°C. The supernatants were transferred to fresh tubes and stored at −20°C for subsequent qPCR analysis.

### Quantitative Polymerase Chain Reaction (qPCR)

qPCR was performed as described previously (Zhang et al. 2016). The primer sequences for measuring target gene transcription levels are listed in **Supplemental Table 5**. The reactions were run in a CFX96 Touch Real-Time PCR Detection System (185-5195, Bio-Rad).

### qPCR Data Analysis

Quantification cycle (Cq) values were calculated by CFX Manager™ software (Bio-Rad). For cDNA qPCR data, the dynamic ranges of RT and amplification efficiency were evaluated before applying the ΔΔCq method to calculate relative gene expression change. The transcription level of the target gene was normalized to the internal control as fold change. For ChIP DNA qPCR data, the Cq value was normalized as percent of input. Data generated in at least three independent experiments are presented as mean ± standard error; the two-tailed Student t-test statistic was applied with P < 0.05 considered to be a significant difference.

### Gene Set Enrichment Analysis (GSEA)

GSEA was performed according to the “GSEA USER GUIDE” (Mootha et al. 2003; Subramanian et al. 2005). For gene ontology (GO) analysis, biological process gene sets from the Molecular Signature Database (MsigDB) were used. Pathways with NOM (Nominal) p-value < 5% and FDR q-value < 25% were included in the analysis.

## Acknowledgements

The authors wish to thank Ms. Diana Kalinowska from KUMC Department of Pharmacology, Toxicology & Therapeutics for her assistance with the ChIP assay. This work was supported by National Institute of Diabetes and Digestive and Kidney Diseases grant R01DK100595 awarded to C. Slawson and K. R. Peterson, R01HL111264 K.R. Peterson, and a KUMC Biomedical Research Training Program grant awarded to Z. Zhang. Partial support was provided by National Institute of General Medical Sciences grants P20GM103549 and P30GM118247.

## Author contributions

Conceptualization: KRP CS.

Data curation: ZZ SG.

Formal analysis: ZZ SG.

Funding acquisition: KRP CS.

Investigation: ZZ SG MPP JDF LVN.

Methodology: ZZ SG DCK KRP MPP JDF LVN CS.

Project administration: KRP CS.

Resources: DCK KRP CS.

Software: ZZ SG.

Supervision: DCK KRP CS.

Validation: ZZ SG MPP JDF LVN.

Visualization: ZZ SG.

Writing: ZZ MPP JDF LVN KRP CS.

